# Transcriptome Profiling of Resistance Genes Analogs in Soybean’s Cross-Tolerance to Water Limitation and Rust Stress

**DOI:** 10.1101/2025.05.20.652543

**Authors:** Gustavo Husein, Thiago Maia, Fernanda R. Castro-Moretti, Jessica D.K. Nunes, Lilian Amorim, Paulo Mazzafera, Harm Nijveen, Claudia B. Monteiro-Vitorello

## Abstract

Asian soybean rust (ASR), caused by *Phakopsora pachyrhizi*, is the most destructive foliar disease of soybean, with yield losses up to 90%. With climate change intensifying drought and expanding disease incidence, it is critical to understand how combined abiotic and biotic stresses influence plant defense. We investigated the transcriptomic response of a susceptible soybean cultivar to ASR infection under normal and water-limited conditions at four infection stages (12, 24, 72, and 192 hours after-inoculation). We observed a biphasic expression of defense-related genes, particularly resistance gene analogs (RGAs), with an early peak at 12 hours and a late resurgence at 192 hours. Combined stress induced a greater number of differentially expressed genes (DEGs) than rust alone, especially at early infection. Among the differentially expressed RGAs (RGADEs), over 64% belonged to the TM-LRR class, and NBS-LRR genes were the most enriched at known ASR resistance loci, particularly Rpp2. Water limitation strongly modulated gene expression at late stages, revealing stress-specific transcriptional reprogramming. These findings reveal cross-tolerance mechanisms in soybean, highlight the temporal dynamics of RGADEs under dual stress, and provide targets for developing cultivars with improved resilience to both rust and water scarcity.

## 1. Introduction

Soybean (*Glycine max (L.) Merr.*) is considered one of the most important crops globally due to its high protein and oil contents, making it a versatile nutritional resource for food, animal feed, and biofuel production (Rahman et al., 2023). The most destructive disease affecting soybeans and the source of severe epidemics is Asian soybean rust (ASR), caused by the fungus *Phakopsora pachyrhizi*. Without chemical control, soybean producers can face productivity losses between 20% and 90%, characterizing the ASR pathogen among the twelve most damaging plant pathogens globally based on its scientific and economic importance (Dean et al., 2012).

Projections of the incidence of *P. pachyrhizi*, considering both the selection pressure on soybeans and climatic changes, point to an increase in disease occurrence in cultivated areas (Alves et al., 2011; Ghini et al., 2007). Soil moisture stress, a significant factor in soybean production losses, is expected to worsen due to climate change (Leng and Hall, 2019). Changes in historical precipitation patterns will likely lead to more severe drought stress in key soybean-growing regions (Thornton et al., 2014). Climate change can further drive the expansion of pathogens and hosts, accelerating the spread of plant diseases to previously unaffected regions (Burdon and Zhan, 2020; Delgado-Baquerizo et al., 2020). Moreover, it can indirectly influence plant-pathogen interactions by altering the biochemical, physiological, ecological, and evolutionary processes of both the host and the pathogen (Cheng et al., 2019; Trivedi et al., 2022; Velásquez et al., 2018). Consequently, the combined effects of biotic and abiotic stresses, such as reduced water availability and pathogen infection, on plant development are being studied more extensively (Camejo et al., 2005; Engelbrecht and Kursar, 2003; Gerós et al., 2016; Mittler, 2006; Palliotti et al., 2009). Additionally, understanding the genetic basis for resistance to these stresses and their interactions remains a critical area of interest (Kakumanu et al., 2012; Le et al., 2012; Xue et al., 2013).

The presence of ASR drastically reduces the photosynthetic capacity of the contaminated leaves. It causes severe defoliation of the plants, effectively reducing the number of pods per plant and the quality and number of seeds (Echeveste Da Rosa, 2015). Fungicides are regularly used to control the fungus. However, in addition to the potentially harmful effects on the environment and the fungicide’s high costs, the low sensitivity of *P. pachyrhizi* to some active ingredients is reducing the availability of effective chemical compounds. Thus, to control ASR, there is a need for more efficient and lasting forms of control (Godoy et al., 2016; Ivancovich et al., 2007; Langenbach et al., 2016).

Among these alternatives, superior soybean varieties with high productivity and resistance to ASR appears to be the most effective way to control the disease (Vuong et al., 2016). To date, soybean cultivars resistant to ASR have been mapped mainly to seven loci, named Rpp 1 to 7, which are specific genomic regions associated with varying degrees of resistance to *P. pachyrhizi* (Childs et al., 2018; Goellner et al., 2010; Kelly et al., 2015; King et al., 2016; Pedley et al., 2019). The loci Rpp1 (Hyten et al., 2007), Rpp4 (Silva et al., 2008), and Rpp6 (Li et al., 2012) are all located on chromosome 18 but at different positions. Additionally, other Rpp loci are found on distinct chromosomes: Rpp2 (Silva et al., 2008) on chromosome 16, Rpp3 (Hyten et al., 2009) on chromosome 6, Rpp5 (Garcia et al., 2008) on chromosome 3, and Rpp7 (Childs et al., 2018) on chromosome 19.

From a functional perspective, immunity has been organized into two layers depending on the cellular response, either by activating extracellular receptors (TM-LRR), also known as pattern recognition receptors (PRR) and usually related to pattern-triggered immunity (PTI), or activating intracellular receptors (NBS-LRR), encoded by disease resistance (R) genes and related to effector-triggered immunity (ETI) (Dodds et al., 2024). These receptors with potential plant resistance activity are collectively termed resistance gene analogs (RGAs) (Sekhwal et al., 2015). Independent of the RGA class, after apoplastic or intracellular perception, the immune response converges to a similar set of downstream events that will potentially prevent infection based on processes such as reactive oxygen species (ROS) production, calcium influx, signaling transduction by mitogen-activated protein kinases (MAPK), defense genes expression, and defense hormones production (DeFalco and Zipfel, 2021).

Next-generation sequencing (NGS) technologies opened opportunities to identify genome-wide, based on sequence similarity, RGAs that encode proteins with structural similarities to *R* genes and their transcription profile when pathogens challenge plants (Rody et al., 2019; Sekhwal et al., 2015). Notably, RGAs often form clusters in plant genomes, which may include functionally related genes that are not necessarily similar in sequence (Chang et al., 2002). This clustering, driven by ancient whole-genome duplications and segmental duplications followed by gene deletions and genomic reorganizations, has expanded RGA families (Michelmore and Meyers, 1998; Perazzolli et al., 2014). Identifying these potential resistance-associated genes and mapping their genomic organization is highly beneficial for plant breeding. This information supports the development of selection strategies that facilitate the early selection of resistant cultivars, reducing costs through approaches such as marker-assisted selection (MAS) (AliFakheri, 2014; Echeveste Da Rosa, 2015).

In a previous report (Castro-Moretti et al., 2024), we observed that water limitation enhanced disease severity caused by *P. pachyrhizi* in soybean, changing significantly leaf metabolic profile. Complementing this previous report, our study aimed to reveal the comparative expression pattern of a susceptible genotype under the influence of reduced water availability in the severity of ASR. Furthermore, we identified RGAs differentially expressed across infection stages under these conditions. Our results will provide a better understanding of key mechanisms involved in disease progression for targeted strategies to develop ASR-resistant soybean cultivars addressing yield stability and global food security.

## 2. Materials and methods

### 2.1. Experimental design, plant material and inoculation

As an obligatory phytopathogen, the *P. pachyrhizi* inoculum was maintained in a greenhouse by regularly inoculating soybean plants. To achieve this, 8 to 10 soybean seeds were sown weekly until they reached the V4 stage. These plants were watered daily until the soil was fully saturated and inoculated with a rust spore solution (10⁵ urediniospores.ml^−1^) in a humid chamber at 23°C in the dark for 24 hours.

The experiment was developed using a fully randomized factorial design, with three biological replicates per treatment of the susceptible soybean commercial cultivar BMX Lança IPRO. These treatments included plants with and without the presence of biotic and abiotic stresses independently, and leaf samples collected at four time points, resulting in the generation of 16 treatments. Soybean plants were kept under two water availability levels: a moderate water deficit, defined as 65% of the plant-available soil water capacity, and a control group of plants cultivated with 80% soil water capacity. The humidity estimation of the soil mixture, prepared in a 2:2:1 (v/v/v) ratio of soil, sand, and manure, at the permanent wilting point was conducted using the procedure detailed by Ferreira (2010). To verify the water status of the treatments, three fresh V2 trifoliate leaves were collected and analyzed for relative water content, following the method described by Barrs and Weatherley (1962).

The plants were grown under controlled temperature conditions in a greenhouse until the development stage V4 for the experimental treatments with controlled water availability. Water control for these plants began 48 hours before inoculation, ensuring that the treatments were well-established. The plants were weighted daily and watered according to their respective water capacity – 80% (control treatment) or 65% (water limited). The plants were then inoculated with the fungus suspension at the same maintenance concentration. Inoculated and mock plants were covered with plastic bags and incubated in the dark in a humid chamber at 23°C (±2°C) for 24 hours. Fungal inoculated and mock plants were maintained separately to avoid cross-contamination of the control plants. After this incubation period, the plants were returned to the greenhouse, where the controlled water conditions were maintained throughout the experiment. One V3 trifoliolate leaf per plant was collected at times 12, 24, 72, and 192 hours after inoculation (HAI) and immediately frozen with liquid nitrogen and stored at -80°C until the extraction of total RNA.

### 2.2. Total RNA extraction and sequencing

Total RNA extraction was performed using the Trizol method (Sigma-Aldrich, St. Louis-MO, USA) and the Purelink RNA mini kit (Invitrogen, Carlsbad-CA, USA), according to the manufacturer’s instructions. After extraction, total RNA was treated with DNAse 1 (Sigma-Aldrich, St. Louis-MO, USA), and RNA quality was verified with electrophoresis on a 1.5% agarose gel and further observation under UV light (2100 Bioanalyzer, Agilent Technologies, Santa Clara, USA). Quantification and purity were verified in a NanoDrop 2000 UV Visible Spectrophotometer (Thermo Scientific, USA).

The RNA library construction by poly-A enrichment was performed using the Illumina TruSeq Stranded mRNA kit (Illumina, Inc., USA), followed by paired-end sequencing using an Illumina NextSeq 2000 with read lengths of 100bp and an average of 10 million clusters or 20 million paired-end reads per sample. In total, 48 independent libraries were sequenced, corresponding to four collection times, two fungal conditions (with and without the presence of the fungal), two water availability regimes (with and without water limitation), and three replicates.

### 2.3. Data processing

The quality of the reads generated in the sequencing was verified through the software SeqKit (version 0.16.1) (Shen et al., 2016). After quality verification, reads were mapped against the Soybean reference genome *Glycine max* (Schmutz et al., 2010) version 4 using the program HISAT2 (version 2.1.0) (Kim et al., 2015). Then, the aligned reads were assembled in their possible transcripts and the gene abundance was estimated using StringTie (version 2.2.1) (Pertea et al., 2015). Subsequently, the prepDE script provided by StringTie was used to generate a count matrix, with genes represented in rows and samples in columns. Genes with a total count of less than ten across all samples were excluded to eliminate low-expression genes and reduce noise from random variation. Expression levels of assembled transcripts summarized to gene level across samples were estimated and compared using the normalization method of DESeq2 (Love et al., 2014).

### 2.4. Gene expression and functional annotation

Principal Component Analysis (PCA) was conducted to explore gene expression patterns using data that was normalized and variance-stabilized transformed (VST) for all genes with the statistical R package DESeq2 (Love et al., 2014). This approach allowed for examining the overall variation in the dataset and visualizing the separation of samples based on the influences of time, biotic stress, and abiotic stress on gene expression. The identification of differentially expressed genes (DEGs) was also performed using DESeq2, considering genes with a False Discovery Rate (FDR) <

### 0.05 as differentially expressed

The over-representation analysis (ORA) of the Gene Ontology (GO) biological process terms enriched in the DEGs was performed using the package topGO (Alexa; Rahnenfuhrer, 2023) from the Bioconductor project (BiocManager v3.18) implementing the Kolmogorov-Smirnov test.

### 2.5. Resistance gene analogs (RGAs)

The RGAs (resistance gene analogs) were analyzed using the encoded proteome of the Soybean reference genome for all different possible gene’s isoforms, based on the characteristics conserved among the resistance genes (*R*) (Hammond-Kosack and Jones, 1997). To this end, six different tools were used to predict resistance genes domains and motifs: 1) Coils v2 (Lupas et al., 1991); 2) TMHMM v2 (Krogh et al., 2001); 3) InterProScan v5.33–72.0 (Zdobnov and Apweiler, 2001); 4) PfamScan with Pfam v32.0 (Bateman et al., 2004); 5) Phobius (Käll et al., 2004); and 6) TargetP 2.0 (Almagro Armenteros et al., 2019).

Each predictor can identify conserved features among the *R* genes by structural analysis. Only sequences containing at least one of the three conserved domains of the RGAs - LRR, NB-ARC, or NB-LRR - were kept to form the set of candidate RGAs for the Soybean reference genome. The RGA candidates were analyzed for their location on the chromosomes and grouped according to (Rody et al., 2019) methodology. Clusters on the genome were formed from at least 3 putative RGAs of any class if there were no more than nine other genes between two RGAs or if the distance between them was less than 250 kb.

While RGA classification was performed at the isoform level due to the potential variability in domain architecture introduced by alternative splicing, downstream expression analyses were conducted at the gene level. In this case, the first annotated isoform for each RGA gene was used as a representative to summarize expression values, including for heatmap visualization.

## 3. Results

### 3.1. Transcriptome Data Analysis

A total of 48 RNA-Seq libraries were generated from leaf tissues of *Glycine max* cv. BMX Lança IPRO, sampled under eight combinations of biotic (Asian soybean rust) and abiotic (water limitation) stress treatments, each applied with and without the respective stress factor, across four time points (12, 24, 72, and 192 hours after inoculation), with three biological replicates per condition. High-quality sequencing reads were obtained from all libraries (**Table S1**). Subsequent quality control and mapping against the *Glycine max* Wm82.a4.v1 reference genome resulted in an average overall read alignment rate of 93.6%. This process identified 86,256 transcripts associated with 52,872 unique genes. After filtering out low-expression genes, 40,012 genes remained, encoding 75.7% of the predicted soybean proteome.

### 3.2. Differential Gene Expression Profiling

Visualization of the first two components of a Principal Component Analysis (PCA) of the samples revealed a predominant clustering effect attributed to the time factor (**Figure 1**). These two components explained 87% of the total variance. Given the predominant clustering by time points, we focused on the specific impact of the other two factors on gene expression. For each time point and per water limitation condition, we compared samples treated with the pathogen against those without the pathogen, resulting in eight pairwise comparisons.

**Figure 1.**
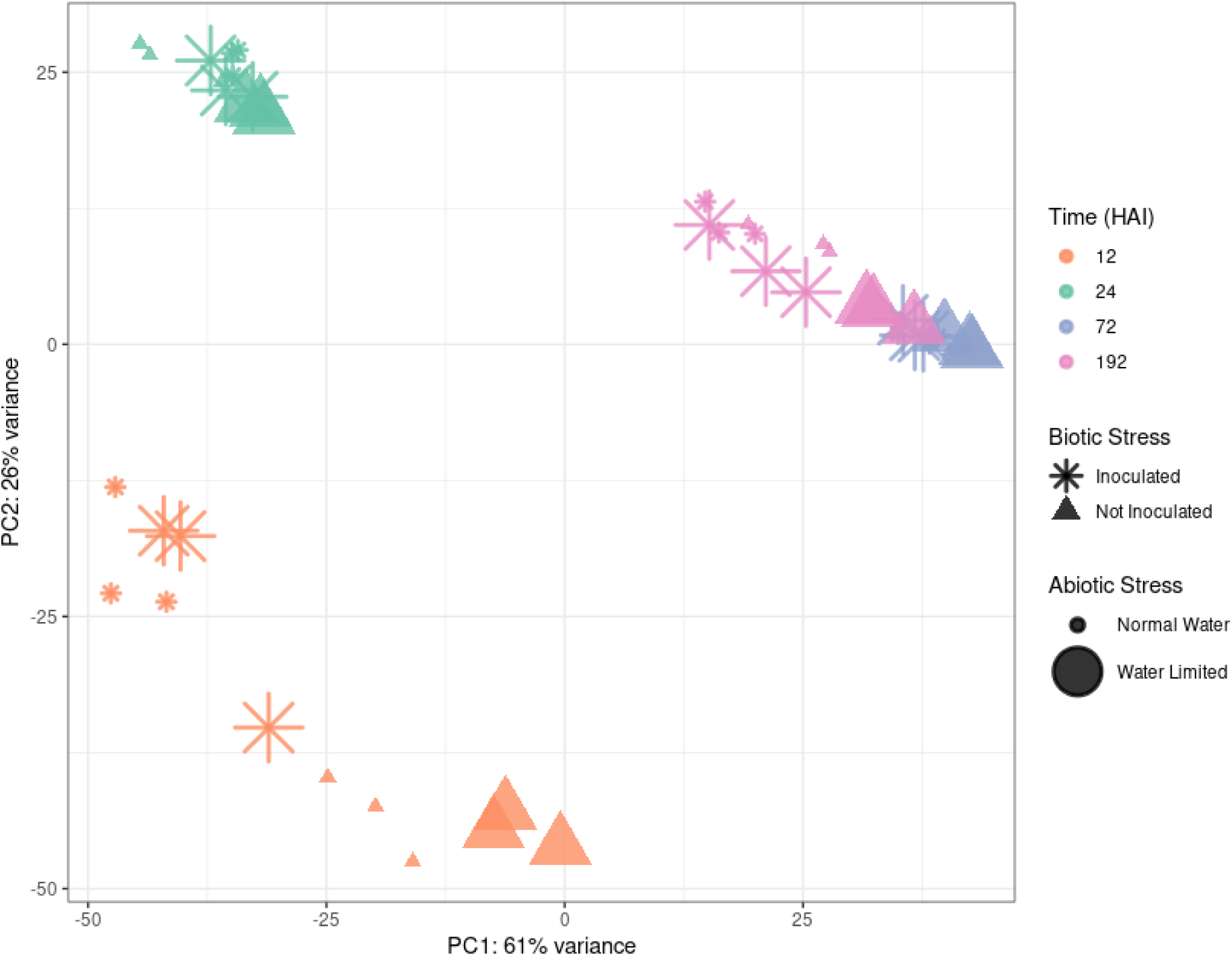
First two components of a Principal component analysis (PCA) of gene expression profiles across time points (12, 24, 72 and 192 HAI) and experimental conditions. Each point represents a biological replicate, colored according to the time point, shaped according to the inoculation treatment (asterisks for inoculated and triangles for non-inoculated samples), and sized according to water availability (smaller symbols for normal water conditions and larger symbols for water-limited conditions).

As a result of this analysis, 9,680, 3,229, 932, and 1,004 differentially expressed genes (DEGs) were obtained for plants in normal water conditions at times 12, 24, 72, and 192 hours after inoculation (HAI), respectively, and 12,666, 1,554, 1,170, and 4,352 DEGs for water-limited plants at the same time points (**Figure 2**, **Table S2**), with almost 65% of the total DEGs obtained from the first time point for both conditions and 33% more DEGs being quantified for water-limited plants.

**Figure 2.**
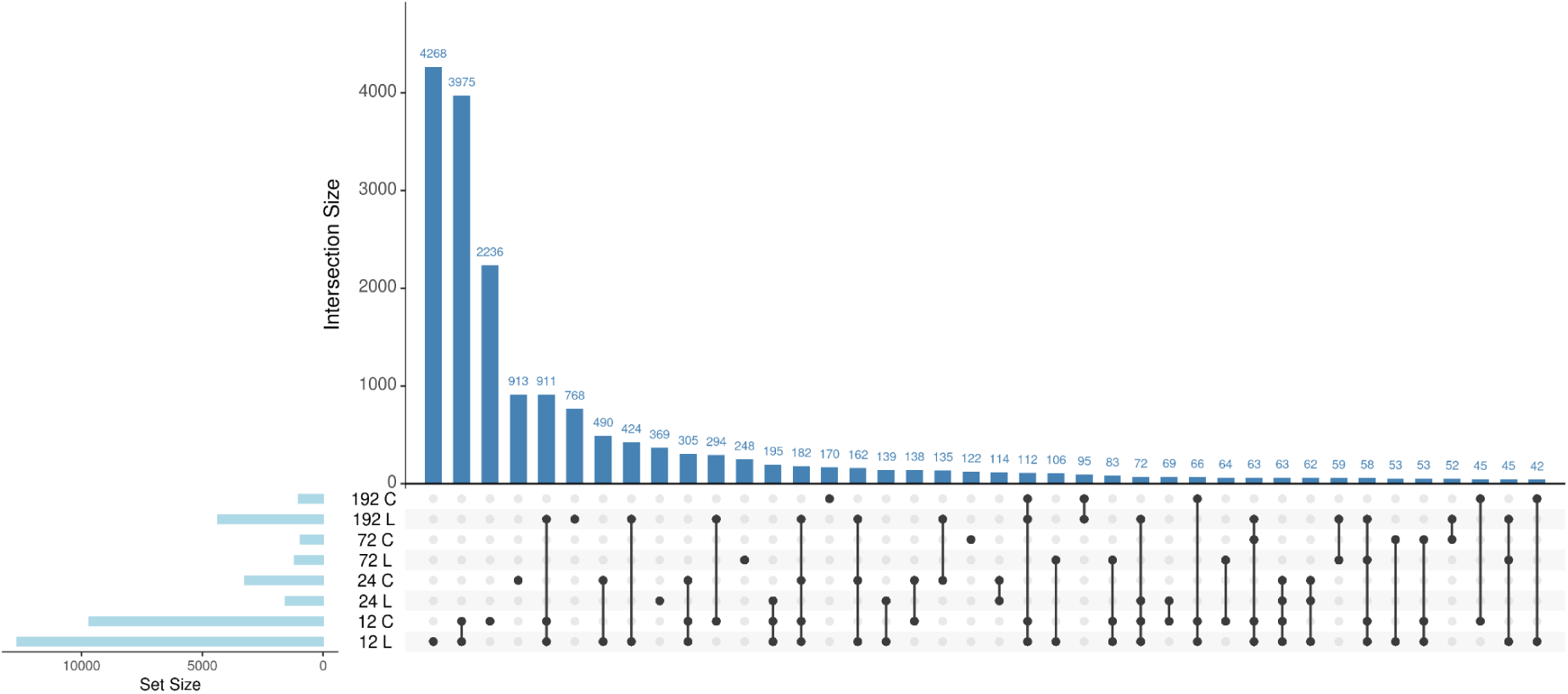
Intersection and number of DEGs across all time points (12, 24, 72 and 192 HAI) and water condition (L: water-limited / C: control). DEGs were identified by comparing pathogen-inoculated samples against non-inoculated samples within each condition and time point, resulting in eight pairwise comparisons. The horizontal bars on the left represent the total number of DEGs identified per condition, while the vertical bars indicate the size of DEG intersections between different conditions. Black dots connected by lines specify the comparisons contributing to each intersection. The numbers above the bars represent the exact intersection sizes.

We also analyzed the overlap between DEGs from different time points and treatments (**Figure 2**) with the objective of identifying shared stress-response genes that may contribute to cross-tolerance mechanisms. The highest number of total DEGs and shared DEGs was observed at the earliest time point (12 HAI), indicating extensive transcriptional changes at this stage across conditions.

### 3.3. Gene Ontology (GO) Enrichment Analysis of DEGs

To identify functional categories potentially enriched in DEGs, we conducted an overrepresentation analysis (ORA) to detect Gene Ontology (GO) terms associated with biological processes in which genes statistically showed differential expression greater than expected by chance. The GO enrichment analysis for the total DEGs among all time points revealed a cumulative total of 2,503 GO terms statistically significant for the control treatment and 2,351 for plants under water limitation. Comparing the two conditions, the number of significant GO terms for control and water limited plants, respectively, were 721 and 644 for 12 HAI, 559 and 512 for 24 HAI, 554 and 561 for 72 HAI and 669 and 634 for 192 HAI (**Table S3**).

Among the top most significant GO terms identified under both water conditions (**Figure S1 and S2**), notable differences emerged across time points regarding genes related to several metabolic processes. At 12 HAI, photosynthesis related terms were prominent in both water-limited and control plants, with an additional enrichment for response to oxidative stress in the control condition, and by 24 HAI, plants under water limitation exhibit enrichment for phenylpropanoid biosynthesis, response to stimulus, and defense response. In contrast, control plants show a broader range of responses, including response to stress, defense response, response to chitin, and response to hypoxia. At 72 HAI, water-limited plants display enrichment for protein phosphorylation, signaling, response to stress and defense response, whereas control plants show enrichment for defense response and response to fungus. Finally, at 192 HAI, water-limited plants highlight processes such as response to stress, defense response, response to biotic and abiotic stimulus, and photosynthesis, while control plants enriched GO terms primarily associated with photosynthesis.

### 3.4. Plant defense-related GO terms

Next, for the plant defense-related GO terms, we calculated the median log_2_ fold change (LFC) of the DEGs (Inoculated vs. Non-Inoculated) per GO term, which represents a quantification of the variation between the expression of genes in plants with the presence of the pathogen compared to plants without the presence of the pathogen. These values were used to demonstrate the variation that the presence of abiotic stress caused in the average expression of genes grouped into biological processes associated with the plant defense response (**Figure 3**). The statistical significance of the differences between the expression levels of control and water stressed plants is provided in **Table S4**.

**Figure 3.**
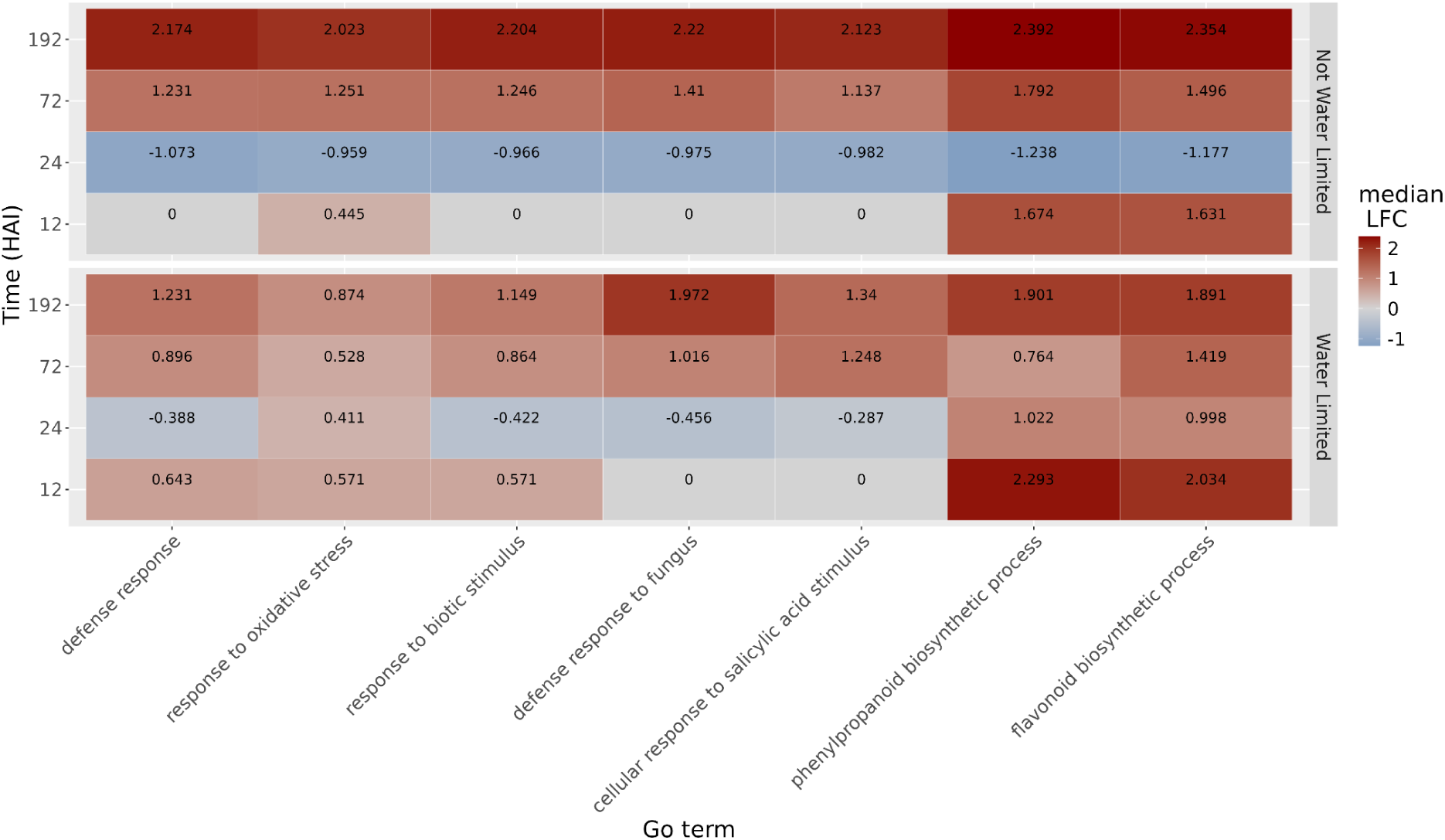
Median log_2_ fold change (LFC) of genes annotated with GO terms related to plant defense response. Non-water-limited plants are represented at the upper part of the graph and the water-limited ones at the bottom. Each row represents a time of collection, while each column represents a GO term associated with plant defense response.

At the beginning of the infection process (12 HAI), upregulated genes are associated with response to oxidative stress, phenylpropanoid biosynthetic process, and flavonoid biosynthetic process, with more pronounced expression changes under water-limited conditions. Concurrently, defense response and response to biotic stimulus were exclusively observed under these conditions. By 24 HAI, the expression levels of these defense-related genes were significantly decreased. In plants under water limitation, defense gene expression returned to basal levels, whereas in control plants, the genes were negatively regulated, showing expression levels below the basal threshold. At 72 HAI, there was a renewed increase in the expression of defense-related genes, particularly notable in plants not subjected to water stress, including genes specifically associated with fungal response, which were positively regulated. This upward trend persisted and intensified until 192 HAI, when defense gene expression remained elevated, especially in plants without water limitation.

### 3.5. Water deprivation effect

To identify the effect of water deprivation on gene expression during infection, DEGs were analyzed between water-limited and control plants, both inoculated with the pathogen. The top 10 most enriched GO terms for each time point are shown in **Figure 4**. At the beginning of the infection process (12 HAI), enrichment is primarily focused on processes related to general metabolic pathways, such as the phenylpropanoid biosynthetic process and secondary metabolite biosynthetic process, alongside stress-related categories such as response to oxidative stress. By 24 HAI, broader stress responses dominate, including terms such as response to organic substance and response to abiotic stimulus. At 72 HAI, terms associated with defense response and response to chemical stimulus remain enriched, reflecting a continuation of general stress responses. However, at 192 HAI, following an extended period of water limitation, a notable shift occurs, with the emergence of genes specifically associated with water-related stress responses, such as response to water deprivation.

**Figure 4.**
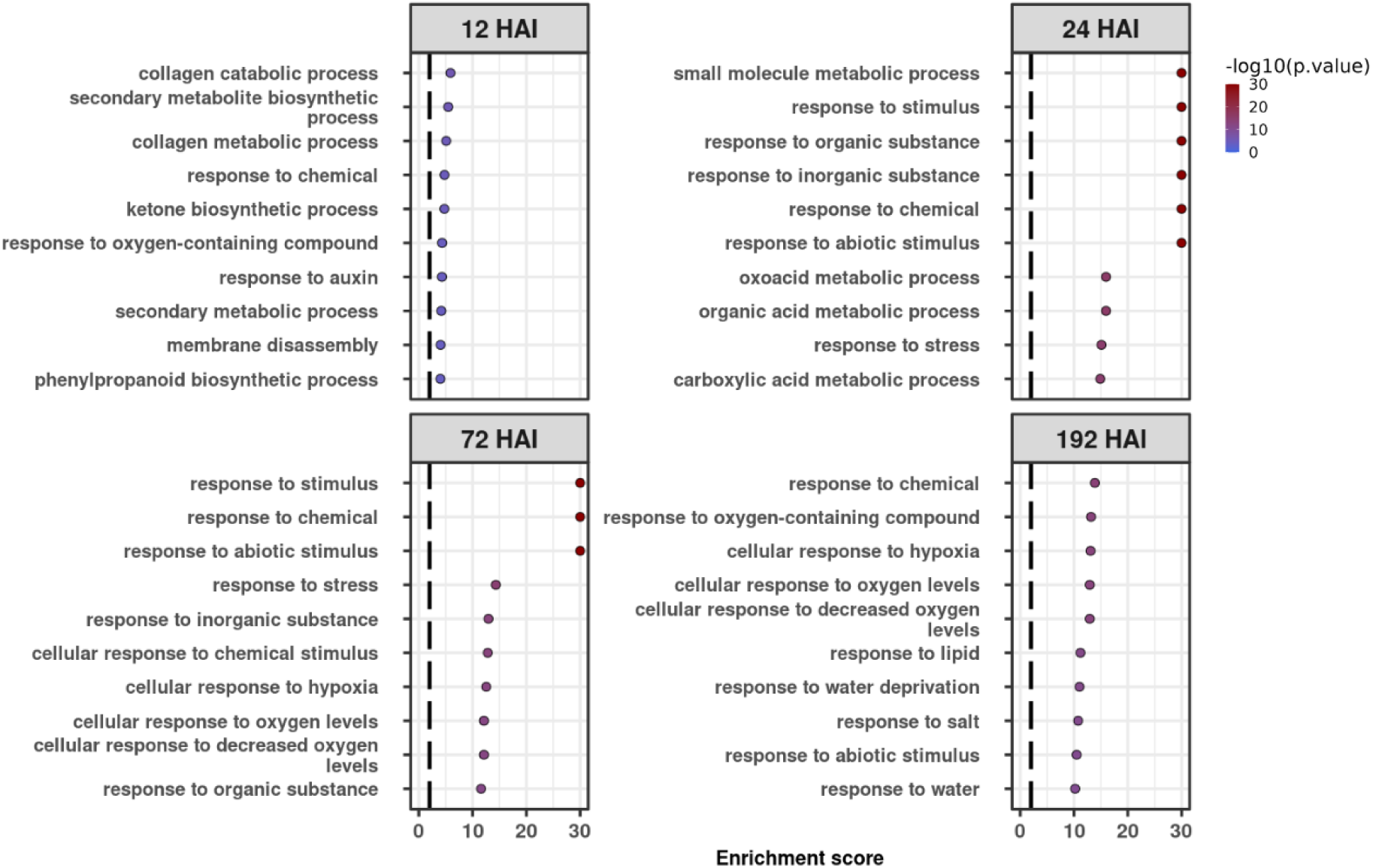
Top 10 enriched Gene Ontology (GO) terms across time points (12, 24, 72 and 192 HAI) for DEGs identified from the comparison between water-limited and control plants inoculated with the pathogen. GO enrichment analysis was performed to identify biological processes (BP) significantly associated with DEGs at each time point. The x-axis represents the enrichment score, while the y-axis lists the enriched GO terms. The color gradient represents the statistical significance (-log_10_(p-value)), with darker shades indicating higher enrichment significance. The vertical dashed line corresponds to a p-value threshold of 0.01.

### 3.6. Prediction of Resistance gene analogs (RGAs)

Using domain prediction tools and a custom Python3 script, the predicted Soybean proteome was scanned for candidate RGA identification, which were subsequently classified based on their combinations of predicted domains (see Material and methods). A total of 4,076 gene isoforms were predicted to encode RGAs classified according to their domain configuration (**Figure 5A**; **Table 1**). Clustering of the predicted RGAs showed that these genes are distributed throughout the genome, with an accumulation of RGAs toward the chromosome ends (**Figure 5A**).

**Figure 5.**
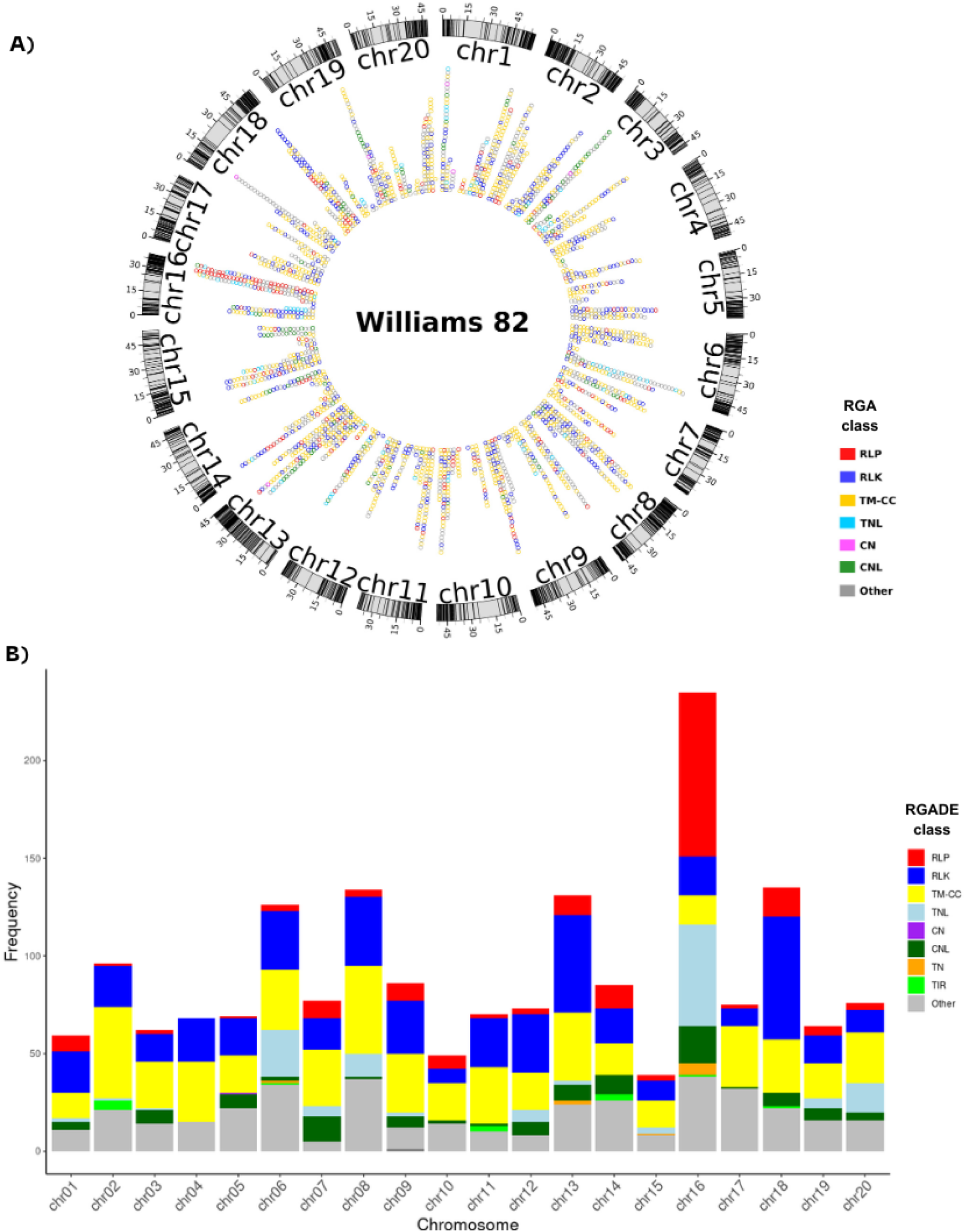
**A)** Distribution of the predicted RGAs in the *Glycine max* genome, mapped across the 20 chromosomes. The predicted RGAs are classified into subclasses, which are represented by different colors. The outer ring indicates chromosome locations in millions of base pairs. **B)** Chromosomal distribution of differentially expressed RGAs (RGADEs) isoforms grouped into subclasses based on their predicted functional domains. The bar heights represent the frequency of each subclass per chromosome, and the colors correspond to the RGA subclasses in (A).

**Table 1.** Number of putative RGAs by domain family and their classes in the reference genome and DEGs (up and down regulated combined).

| RGA class | Whole genome | Overall DEGs | Water Limited DEGs | Control DEGs |
| --- | --- | --- | --- | --- |
| NBS-LRR encoding |  |  |  |  |
| CNL | 264 (6.48%) | 106 (5.80%) | 58 (4.00%) | 86 (7.26%) |
| TNL | 225 (5.52%) | 130 (7.11%) | 115 (7.92%) | 43 (3.63%) |
| TN | 26 (0.64%) | 10 (0.55%) | 10 (0.69%) | 7 (0.59%) |
| CN | 9 (0.22%) | 3 (0.16%) | 3 (0.20%) | 3 (0.25%) |
| TM-LRR encoding |  |  |  |  |
| RLK | 916 (22.47%) | 464 (25.40%) | 372 (25.64%) | 345 (29.14%) |
| RLP | 340 (8.34%) | 189 (10.35%) | 149 (10.27%) | 126 (10.64%) |
| Other variants |  |  |  |  |
| TM-CC | 1158 (22.41%) | 518 (28.35%) | 416 (28.67%) | 337 (28.46%) |
| TL | 1 (0.02%) | 1 (0.05%) | 0 (0%) | 1 (0.08%) |
| TIR | 50 (1.23%) | 14 (0.76%) | 14 (0.96%) | 8 (0.68%) |
| Other | 1087 (26.67%) | 392 (21.45%) | 314 (21.64%) | 228 (19.25%) |
| Total Number of RGAs | 4076 (100%) | 1827 (100%) | 1451 (100%) | 1184 (100%) |

Five classes of RGAs were predicted most frequently, CNL: coiled coil (CC) associated with NB-ARC and leucine-rich repeats (LRR); TNL: TIR domain associated with NB-ARC and LRR; RLK: receptor-like kinase; RLP: receptor-like protein; and TM-CC: transmembrane domain associated with a coiled-coil domain (**Table 1**). Furthermore, differentially expressed RGA (RGADE) isoforms were observed across all chromosomes with a higher number on chromosomes 2, 6, 8, 13, 16 and 18, accounting for almost 50% of the total RGAs predicted (**Figure 5B**; **Table 1**). The most frequently predicted RGA classes were those containing a transmembrane domain, including RLK and RLP (TM-LRR encoding class), as well as the TM-CC subclass. Considering the total number of RGAs predicted in the genome, they correspond to 53%, and among DEGs, they represent 64% of all expressed RGAs combining both water conditions. This tendency further intensifies when DEGs are separated based on water limitation, reaching 68% for plants under normal water condition. On the other hand, in conditions with only biotic stress, the proportion of these RGAs aligns more closely with the overall set of RGADEs, maintaining around 64%.

When assessing the effect of water limitation, we observed notable variations in RGADE expression between plants exposed to water limitation and those maintained under controlled conditions, when compared to the reference genome. For instance, the TNL subclass exhibited expression exclusively under water limitation; isoforms from the RLK subclass were expressed in both conditions but at a higher level in control plants, and the RLP subclass showed a more moderate expression profile.

The presence of the pathogen affected the proportion of RGAs relative to the total number of gene isoforms. In the reference genome, 4,076 out of 66,210 gene isoforms (6.15%) were classified as RGAs. This proportion was higher among DEGs obtained by comparing inoculated versus non-inoculated plants, with 1,827 out of 19,165 DEGs isoforms (9.53%) identified as RGAs. Under water-limited conditions, 1,451 out of 15,452 isoforms (9.39%) were classified as RGAs, compared to 1,184 out of 12,653 isoforms (9.36%) under control conditions. These results demonstrate an enrichment of RGAs among DEGs driven by pathogen inoculation, with no substantial impact from water stress on the proportion of RGAs among expressed genes.

The timing of RGADE expression largely coincides with early pathogen invasion, particularly at 12 HAI, when tissue penetration is critical. Under water limitation, 1074 (56.76%) of RGADEs were expressed at 12 HAI, followed by a sharp drop to 112 (5.91%) at 24 HAI, 136 (7.18%) at 72 HAI, and 570 (30.12%) at 192 HAI. In contrast, under non-limiting water conditions, 788 (55.85%) of RGADEs were expressed at 12 HAI, 313 (22.18%) at 24 HAI, 203 (14.38%) at 72 HAI, and only 107 (7.58%) at 192 HAI (**Figure S3; Table S5**). These results suggest that water availability modulates the temporal distribution of RGADE expression, particularly at later infection stages.

### 3.7. Expression profile of resistance gene analogs differentially expressed

After exploring RGA isoform-specific expression, we collectively examined the expression profiles of the 932 RGADEs at the gene level (**Figure S4**). We observed a distinct temporal expression pattern between plants exposed to limited water and control plants. Under water-limited conditions, 54.85% of RGADEs were identified at 12 HAI, followed by 6.77% at 24 HAI, 8.48% at 72 HAI, and 29.9% at 192 HAI. In contrast, control plants exhibited a markedly different distribution, with 50.71% of RGADEs expressed at 12 HAI, 24.15% at 24 HAI, 15.96% at 72 HAI, and only 9.18% at 192 HAI. These results highlight a pronounced enrichment of RGADE expression at 192 HAI under water-limited conditions, while control plants showed a more uniform distribution of RGADEs across earlier time points, particularly at 24 and 72 HAI.

Of the RGAs identified within the Rpp loci (55 genes), 28 were differentially expressed (RGADEs). These RGADEs were distributed across most loci and treatments, except for Rpp7 (**Figure 6**). At 12 HAI, the water limitation resulted in the induction of RGADEs of classes TNLs (6), RLK (1), and Other (3), whereas control plants downregulated RGADEs of classes TM-CC (1) and TNLs (4). At 24 HAI, RGADEs were nearly absent in water-limited plants, while control plants displayed a diverse expression pattern, including both upregulated (e.g., 2 Other and 5 TNLs) and downregulated (e.g., 3 Other, 1 TN, and 1 CNL) RGADEs. At 72 HAI, both conditions resulted in upregulated genes in low numbers. By 192 HAI, plants exposed to water stress showed exclusively upregulated RGADEs of most classes: TNLs (4), RLP (1), Other (3), TM-CC (1), and TN (1), while control plants had only one upregulated TM-CC.

**Figure 6.**
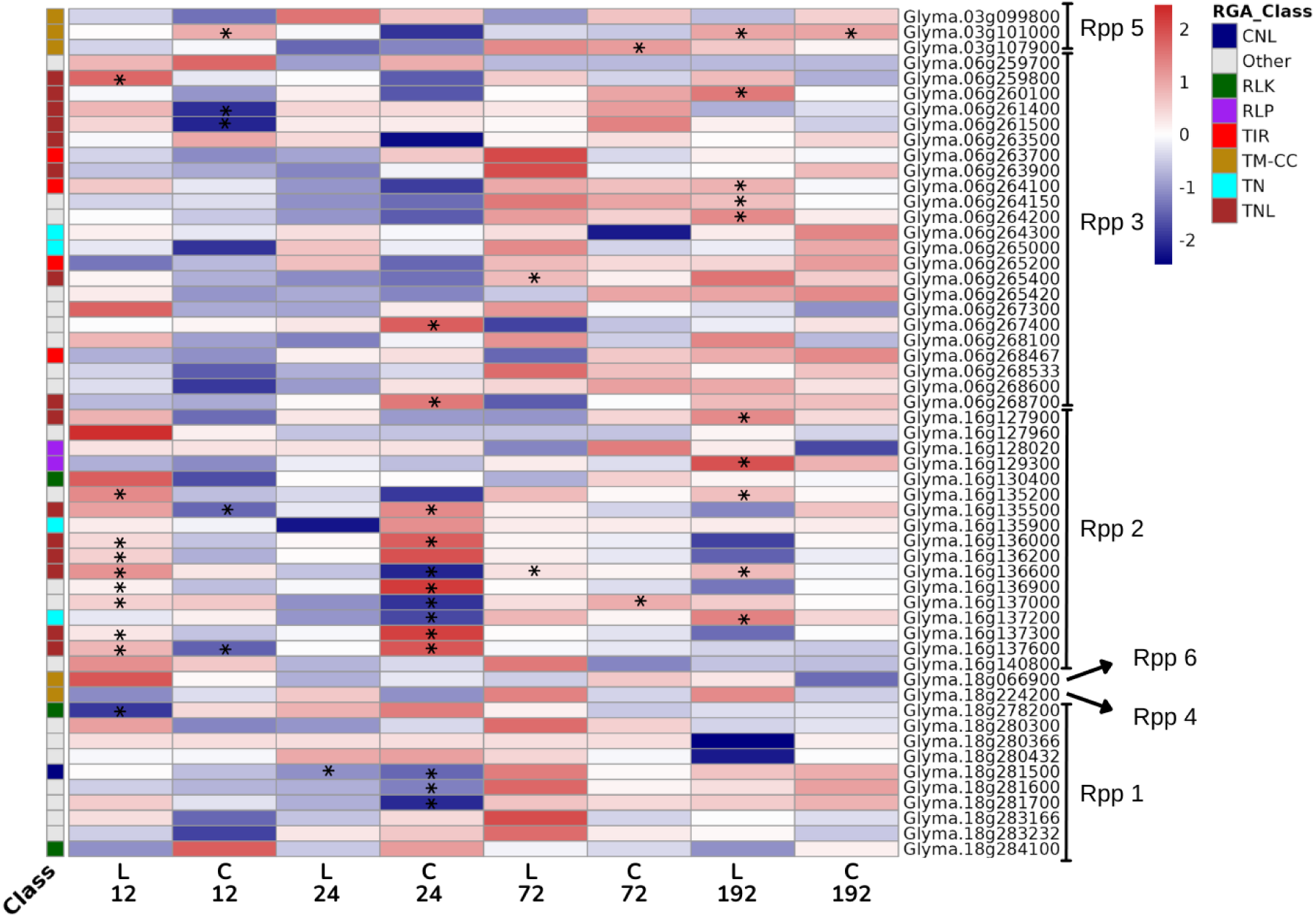
Heatmap of RGAs expression at Rpp loci across all time points (12, 24, 72 and 192 HAI) and both water conditions (L: water-limited / C: control). The color scale indicates the log_2_ fold change for inoculated vs. not-inoculated plants, with red representing upregulation and blue representing downregulation. The RGA subclass of each gene is indicated by the color-coded bar on the left. Asterisks (*) denote DEGs. Annotations on the right indicate Rpp loci (Rpp1, Rpp2, Rpp3, Rpp4, Rpp5, and Rpp6) where these RGAs are located.

Notably, the loci Rpp2 (12 RGADEs, 43%) and Rpp3 (10 RGADEs, 35.7%) grouped the highest number of genes modulated in our experimental conditions (**Figure 6**). Respectively, the analysis of loci Rpp5 and Rpp1 revealed only 2 (7.1%) and 4 (14.3%) RGADEs. In addition, among all treatments, the RGADEs classified as TNLs were the most represented (52%).

## 4. Discussion

The interaction between climate-induced changes in global water availability and the susceptibility for plant diseases such as Asian Soybean Rust (ASR) remains unpredictable and critically important for the future of crop breeding (Jorge et al., 2015; Konapala et al., 2020). For some time, various authors have been exploring the crosstalk between biotic and abiotic stresses, considering their effect as either positive or negative in different pathosystems (Choudhary and Senthil-Kumar, 2024; Singh et al., 2023; Sunarti et al., 2022). Also, we cannot assume that individual responses to different stresses can predict the effects of combined stresses (Pandey et al., 2015). As already demonstrated in diverse pathosystems (Paoletti et al., 2001; Ragazzi et al., 1995; van NIEKERK et al., 2011), water deficit imposes a significant drought-stress penalty on plants, which has been widely associated with the aggravation of fungal diseases (Swett, 2020) and negative impacts on crop development (Ghosh and Roychoudhury, 2024). The effects of drought facilitate pathogen colonization, increasing disease incidence, and exacerbating symptom severity due to the suppression of the plant defense mechanisms (Boyer, 1995; Swett, 2020). These findings underscore the critical role of water availability in maintaining effective plant defenses against fungal pathogens.

In this context, we examined the impact of water limitation on the transcriptomic response of a susceptible soybean genotype to *P. pachyrhizi*, analyzing infected leaves at four specific time points. The time points chosen represent key phases of the fungal infection process, including the formation of appressoria, cuticle penetration, and the invasion and subsequent growth of hyphae within the host tissue, observed at 12, 24, 72, and 192 hours post-infection, respectively (Gupta et al., 2023).

The first overall analysis revealed that time has the most profound effect on the transcriptome, irrespective of the inoculation state. The pairwise comparison per time point of inoculated plants vs. inoculated plants under limited water conditions showed that genes related to plant defense (GO:0006952) had an intensified response in plants under water limitation at the early stages of infection. This effect diminishes as time passes and disease progresses. The number of differentially expressed genes showed a biphasic response, as seen before with the same pathosystem (van de Mortel et al., 2007), with defense-related genes being upregulated at earlier stages of infection (12 HAI), followed by a down-regulation (24 HAI) and then again being positively regulated at later time points (72 and 192 HAI).

We unveiled 33% more DEGs in plants with combined stress than in plants exposed to only the biotic stress. This higher number of DEGs reflects the broader range of plant responses necessary to maintain physiological homeostasis, coordinated by gene expression (Pandey et al., 2015; Zandalinas et al., 2018). Among the DEGs identified at 12 HAI, the strongest upregulated under water limitation were *Glyma.18g211100* (Peroxidase), *Glyma.18g267900* (Isoflavone7-O-methyltransferase-like protein), *Glyma.02g007400* (Chitinase) and *Glyma.14g205200* (Cytochrome P450). Their expression levels were between 7 and 86 times higher in inoculated plants compared to non-inoculated plants under water limitation. The induction of these genes may lead to the production of phytoalexin, lignin, flavonoids, ABA, and chitin-pathogen digestion, which serve as chemical and physical barriers against fungal infection. These immune system components may also play roles in the response to water limitation (Almagro et al., 2009; Li et al., 2022; Zhang et al., 2017). Moreover, these genes are involved in other stress responses in soybean plants. Functional characterization of *Glyma.18g211100* has been shown to contribute to resistance against Cercospora leaf blight (Patel et al., 2024), *Glyma.18g267900* and *Glyma.02g007400* are associated with resistance to soybean cyst nematode (Hu et al., 2024; Zhang et al., 2017), while *Glyma.14g205200* was associated with drought resistance (Li et al., 2022). The plant’s adaptability to diverse environmental conditions depends on a versatile defense network of genes with sometimes overlapping functions, which we also detected in our work.

The robust gene expression of plants exposed to combined stresses may stimulate transcriptional changes, leading to a faster and more intense response upon exposure to subsequent stress, as described for the priming effect (Mauch-Mani et al., 2017). Indeed, our data showed 25% more RGADE isoforms in plants exposed to combined stresses compared to only rust-inoculated plants. Additional evidence of the effects of combined stresses was indicated by the increased expression of RGAs in the early stages of infection, suggesting the activation of stress defense mechanisms under these conditions. Approximately 36% of the total RGADEs encoded receptor-like kinases (RLKs) and receptor-like proteins (RLPs). Considering that TM-LRR proteins constitute 30% of all RGAs in the genome, our analysis revealed an enrichment of this receptor class under both experimental conditions—ASR inoculation with and without water limitation.

These proteins include pattern recognition receptors (PRRs), which recognize microbial-associated molecular patterns (MAMPs), resulting in pattern-triggered immunity (PTI) (Jones and Dangl, 2006). Some PRRs may also function as sensors of developmental changes due to abiotic stresses, which could explain the increased number of RGADEs detected in the combined stresses. Notably, an increased abundance of RGADEs encoding RLKs was observed in water-limited plants compared to controls at 192 HAI, a time point linked to the plant’s response to prolonged water scarcity. This finding aligns with the dual role of RLKs in mediating responses to both biotic and abiotic stresses, as they are known to play a critical role in osmotic stress signaling in various plant species (Osakabe et al., 2013).

Therefore, we propose that water limitation enhances the plant’s perception of multiple stressors, activating similar receptor types and leading to a stronger PTI response in susceptible plants. Despite the activation of these receptors, the overall diminished defense response observed in water-limited plants at later stages, when the fungus has already established itself and proliferated within the plant, suggests that effector-triggered susceptibility (ETS) significantly contributes to disease progression and severity.

Throughout their infection stages, biotrophic fungi overcome plant defense systems by directing various effectors to impair plant defenses, whether suppressing PTI-activated responses after pathogen recognition or interacting with ETI receptors (Toruño et al., 2016). ASR fungus presents an efficient strategy to defeat responses upon that perception, leading to a strong downregulation of genes related to plant defense at 24 HAI. The low representation of NBS-LRR-encoding genes among RGADEs at this time suggests a potential ETI response deactivation related to the expression and release of fungal effectors proteins (Gupta et al., 2023). Other authors identified similar patterns for rusts, considering, for instance, time after inoculation for the analysis (Dobon et al., 2016; Pradeu et al., 2024). The reduced expression of defense genes in plants under water stress at 24 HAI is not as marked but still noticeable. Following this wave down, an increment toward positive net expression reaches its maximum at 192 HAI. With disease progress and fungal multiplication, the increase in MAMPs may result in a new wave of pathogen perception, reactivating the innate response at later moments. At that point, the induction of genes in which GO terms refer to plant defense, such as response to chitin and oxidative stress, has no effect since the disease has already been established and sporogenesis initiated.

The lower impact of the waves of recognition in water-limited plants, along with an increased expression of genes related to response to water deprivation, suggests a redirection of efforts otherwise used for controlling the pathogen’s development, resulting in a more robust plant susceptibility (Beattie, 2011; Leisner et al., 2023).

We also analyzed the distribution of RGAs across chromosomes and observed that RGAs tend to accumulate at the chromosomal ends, organizing gene clusters. This distribution is not uncommon, as has been described for RGAs in other crop’s genomes, and is also similar to the distribution of protein-coding genes in the soybean genome (Christie et al., 2016; Liu et al., 2020; Rody et al., 2019; Wang et al., 2021). Additionally, for soybean, seven loci harboring a combination of various RGAs, conferring race-specific resistance to a limited number of *P. pachyrhizi* isolates, were identified and genetically mapped to the soybean genome (Rpp 1 to Rpp 7) (Childs et al., 2018). As described, only Rpp7 did not include RGADEs in our experiments. All the others harbor RGADEs (50% of all RGAs), with eleven genes shared between water-limited and control conditions at specific time points. Notably, nine of these RGADEs were within the Rpp2 locus encoding disease resistance proteins of the TIR-NBS-LRR (TNLs) class. TNLs are active components of the ETI response, which seems suppressed in our experiments. We speculate that Rpp2 is the most relevant locus for the *P. pachyrhizi* infection in the susceptible genotype used in our experiments, as the earlier induction of these genes is in water-stressed plants.

The Rpp2 locus offers additional insights into their coordinated expression and functional significance during pathogen attack. Among the RGAs, *Glyma.16g136000*, *Glyma.16g136900*, *Glyma.16g137000*, *Glyma.16g137300,* and *Glyma.16g137600* were all induced at 12 HAI under water-limited conditions. However, Glyma.16g136600 was consistently induced in water-limited plants but repressed 24 HAI in regularly watered plants, suggesting a role in abiotic stress. This is an intricate network that ties the clustering of genes and possibly co-regulation to enable a rapid and localized immune response. The clustering of resistance genes is a common feature in plant genomes derived from selective pressures that favor the retention and duplication of genes involved in pathogen recognition and defense (Shao et al., 2016). However, despite the upregulation of these closely linked RGAs, particularly under water-limited conditions, the pathogen continues to establish and proliferate within the host.

We also identified DEGs related to abiotic stress to further weaken the effectiveness of a coordinated response toward combined stresses. For instance, the antagonistic effect of abscisic acid (ABA) on salicylic acid (SA)-mediated pathways. SA is crucial for activating defense responses against biotrophic pathogens (Yasuda et al., 2008). ABA is a key hormone driving plant responses to drought by regulating stomatal closure, osmotic balance, and other mechanisms critical for drought adaptation. The ABA-SA interaction may suppress the expression of pathogenesis-related genes, compromising the plant’s immune system (Ghosh and Roychoudhury, 2024). As a result, the activation of ABA pathways under drought stress may prioritize abiotic stress adaptation at the expense of biotic stress resistance, thereby facilitating pathogen establishment and proliferation.

Exploring the regulatory networks governing RGA cluster modulation and their interactions with abiotic stress pathways is essential to better understanding the constraints on effective immunity in soybean genotypes. Our study emphasizes the significant modulation of soybean immune responses by water availability and identifies candidate genes for further investigation. These findings enhance our understanding of the complex interplay between abiotic and biotic stress responses, guiding the development of comprehensive strategies to manage drought and disease.

## Acknowledgments

Dr. Quirijn de Jong van Lier (Soil Physics Laboratory, CENA/USP) for helping us to estimate the soil humidity at the permanent wilting point. Carlos Martinelli and GDM Seeds for their generous donations of soybean seeds. Dr. Sergio Pascholati and Ms. Sabrina Holz for sharing their *Phakopsora pachyrhizi* population. CBMV, LA and PM thank CNPq (The Brazilian National Council for Scientific and Technological Development) for research fellowships. GH was supported by Coordenação de Aperfeiçoamento de Pessoal de Nível Superior (CAPES 88887.671506/2022-00). This study was supported by the Fundação de Amparo à Pesquisa do Estado de São Paulo (FAPESP – 2019/13191-5).

## Conflict of Interest statement

The authors declare that the research was conducted in the absence of any commercial or financial relationships that could be construed as a potential conflict of interest.

## Notes

### Competing Interest Statement

The authors have declared no competing interest.

