## Supplementary figures and images for "Transcriptome Profiling of Resistance Genes Analogs in Soybean’s Cross-Tolerance to Water Limitation and Rust Stress"

### Supplementary Figure 1

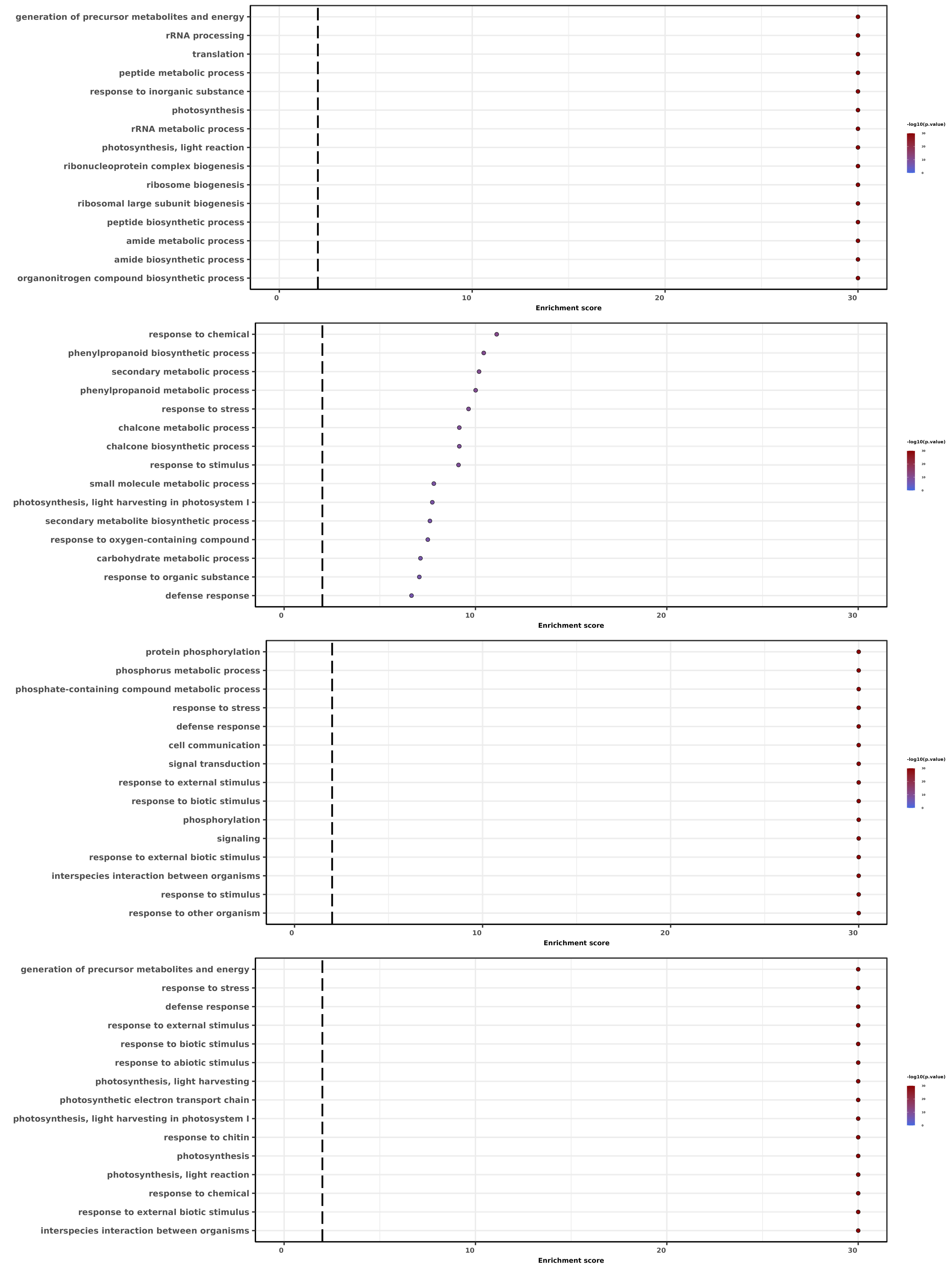

### Supplementary Figure 2

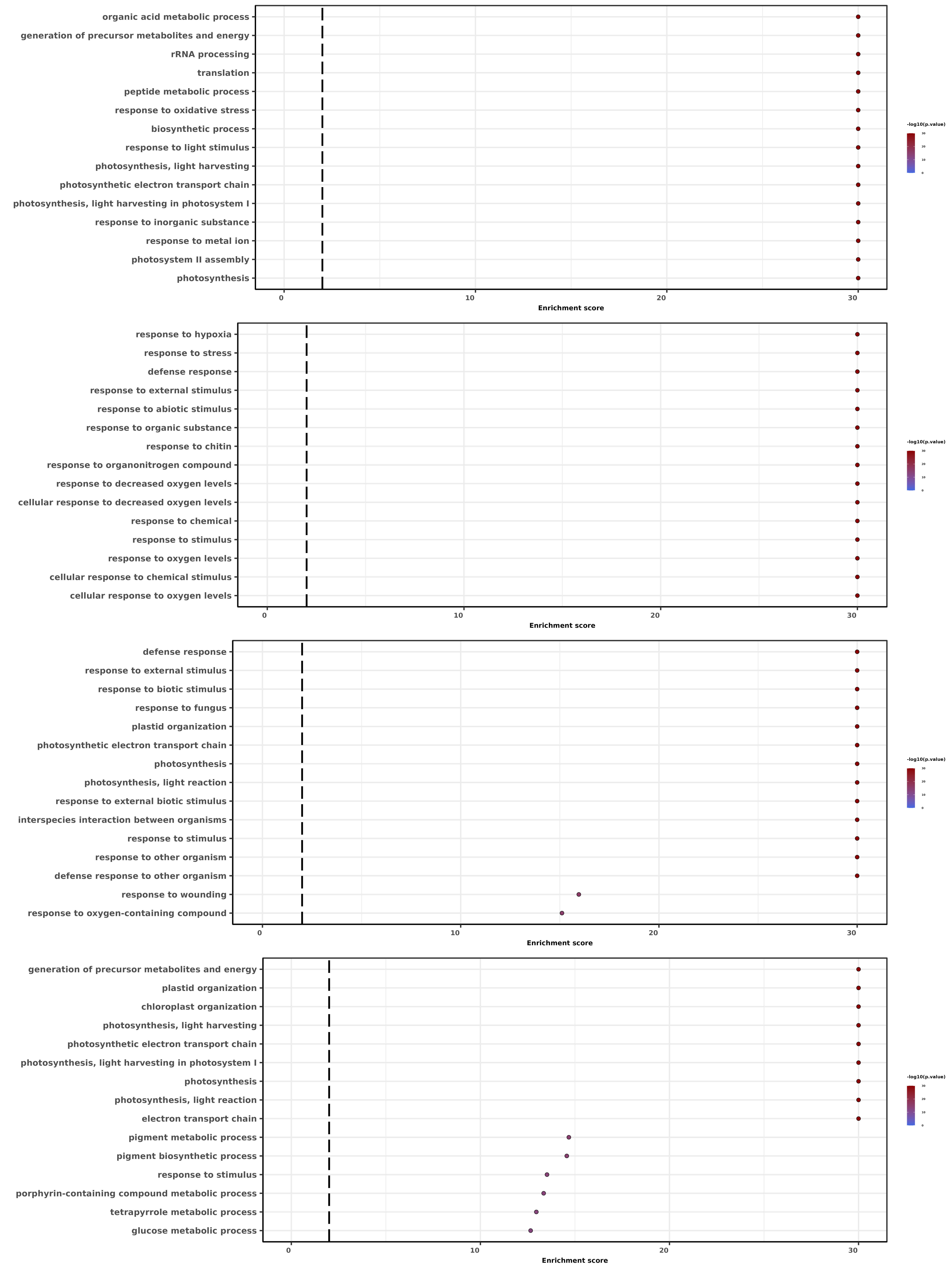

### Supplementary Figure 3

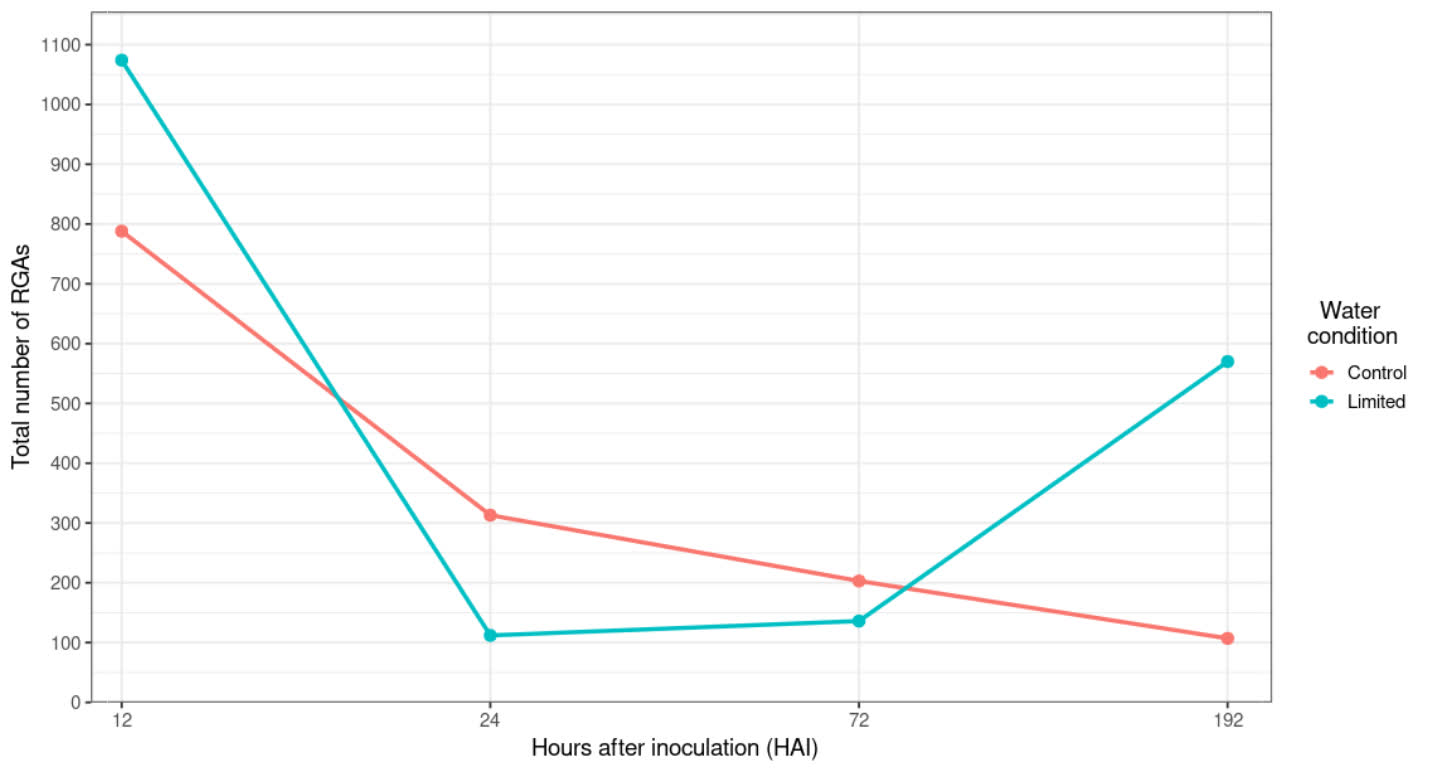

### Supplementary Figure 4

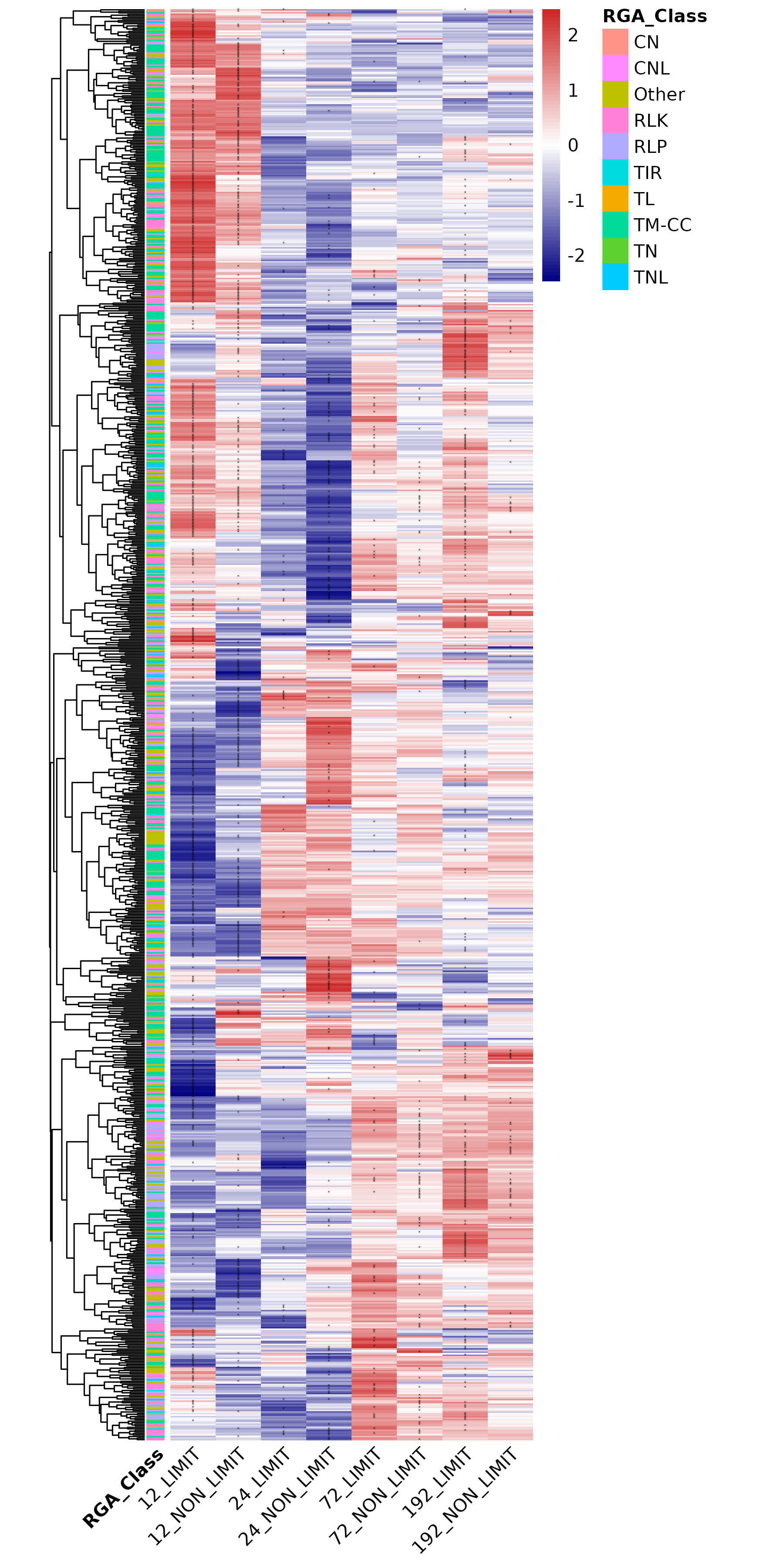
